# The first steps toward a global pandemic: Reconstructing the demographic history of parasite host switches in its native range

**DOI:** 10.1101/2020.07.30.228320

**Authors:** Maeva A. Techer, John M. K. Roberts, Reed A Cartwright, Alexander S. Mikheyev

**Author notes:** Texas A&M University, Department of Entomology, Biological Control Facility, College Station, Texas 77840, USA. **Corresponding Authors:** Maeva A. Techer, Alexander S. Mikheyev.

## Abstract

**Background:** Host switching allows parasites to expand their niches. However, successful switching may require suites of adaptations and also may decrease performance on the old host. As a result, reductions in gene flow accompany many host switches, driving speciation. Because host switches tend to be rapid, it is difficult to study them in real-time and their demographic parameters remain poorly understood. As a result, fundamental factors that control subsequent parasite evolution, such as the size of the switching population or the extent of immigration from the original host, remain largely unknown. To shed light on the host switching process, we explored how host switches occur in independent host shifts by two ectoparasitic honey bee mites (*Varroa destructor* and *V. jacobsoni*).

**Results:** Both switched to the western honey bee (*Apis mellifera*) after it was brought into contact with their ancestral host (*Apis cerana*), ∼70 and ∼12 years ago, respectively. *Varroa destructor* subsequently caused worldwide collapses of honey bee populations. Using whole-genome sequencing on 63 mites collected in their native ranges from both the ancestral and novel hosts, we were able to reconstruct the known temporal dynamics of the switch. We further found multiple previously undiscovered mitochondrial lineages on the novel host, along with the genetic equivalent of tens of individuals that were involved in the initial host switch. Despite being greatly reduced, some gene flow remains between mites adapted to different hosts.

**Conclusion:** Our findings suggest that while reproductive isolation may facilitate the fixation of traits beneficial for exploitation of the new host, ongoing genetic exchange may allow genetic amelioration of inbreeding effects.

## Background

Arms races between parasites and their hosts drive evolutionary innovation. Novel parasites can decimate host populations or drive them extinct unless counter-adaptations evolve. Similarly, parasite evolution accompanies the very act of host switching since it requires adaptations to novel host physiology and behaviour to persist and to spread. Because parasite adaptations tend to be host-specific, host switches are often associated with host-associated genetic differentiation and eventual speciation. However, only the endpoint of this process is typically observed, as host switches tend to occur rapidly, and the original host is often unknown. As a result, many unanswered questions remain about how parasites acquire new hosts. For instance, if host switches are accompanied by a bottleneck due to reduced gene flow from the ancestral host, how does the parasite have sufficient genetic diversity to adapt? Does gene flow cease completely, or does it continue at a low level, potentially providing additional genetic material for adaptations?

One of the major limiting factors for host switching is the geographic separation between parasites and potential hosts [1]. Globalization has eased these barriers, sometimes giving rise to pandemics [2]. As a result, host switches are easier to observe and to study in something approaching real-time. One of the most dramatic and economically important switches involved the two ectoparasitic mite species, *Varroa destructor* recently followed by *Varroa jacobsoni*, which acquired the western honey bee (*Apis mellifera*) as a new host, ∼70 and ∼12 years ago, respectively [3,4]. *V. destructor,* in particular, spread worldwide, causing extensive honey bee population collapses, whereas *V. jacobsoni* has so far remained in Oceania [5,6]. Both mites were originally found on the sister species, *Apis cerana*, and came into contact with growing populations of *A. mellifera,* which were brought in for purposes of beekeeping [7,8]. These two host switches occurred in parallel and relatively recently, allowing the reconstruction of how the host switches took place using genomic tools.

Both switches have been investigated using microsatellite markers and relatively short mitochondrial markers, which revealed that in both species populations on the new host were strongly differentiated and genetically depauperate [4,9–13]. While *V. destructor*, in particular, was described as “quasi-clonal” and highly inbred [9], it successfully spread worldwide and has shown a remarkable ability to parasitize genetically diverse *A. mellifera* populations, as well as to evolve resistance to human counter-measures, such as synthetic acaricides How does a bottlenecked species with high inbreeding achieve such a level of success? Increasing evidence from population genomic analysis of fungal pathogens suggests that the success of many pathogens appears to rely on maintaining some level of adaptive diversity despite the presence of bottlenecks during host switches [14,15] This is also true of many invasive animals who overcome genetic bottlenecks with repeated introductions or other mechanisms such as multiple mating [16–19], making the case of *V. destructor* that much more puzzling.

To answer this question and to gain broader insight into how host switches happen, we sequenced nuclear and mitochondrial genomes from sympatric populations of the two mites across Asia and Oceania, collected on both novel and introduced hosts. This allowed us far greater power to examine how the host switch took place with much greater precision than was possible previously. We found strikingly parallel dynamics at play in both host switches, which were characterized by a surprisingly large effective population size at the time of the switch and ongoing gene flow with cryptic population genetic processes that may have helped *Varroa spp.* succeed.

## Results

### Cryptic diversity in mitogenomes suggests multiple foundresses at the origins of the host switch

We examined the genome-wide variation and divergence among 31 *V. destructor* and 32 *V. jacobsoni* females from their original and novel hosts (Table 1). *Varroa* species identity was confirmed by extracting and aligning mitogenomes together with known reference sequences of the mtDNA *COX1* 458-bp standard marker (Table S1 and Figure S1). A sole mismatch between taxonomic identification and mtDNA barcoding was detected in a *V. jacobsoni* specimen (THV003_1), which was collected in Thailand on *A. mellifera,* though it is not known whether the mite could reproduce on this honey bee species.

**Table 1:** Varroa spp. specimens used for population genomics were collected from their native range across 11 countries, both on their original or new honey bee host. Newly reported mtDNA lineages are indicated in bold.

| Species | Host | Country | Date | N | mtDNA lineages as defined by standard *COX1* |
| --- | --- | --- | --- | --- | --- |
| *V. destructor* | Original host<br>*A. cerana* | Japan | 1994 | 2 | VD Japan J1 |
|  |  | South Korea | 1996 | 1 | VD Korea K1 |
|  |  | China | 2001-2002 | 6 | VD China C1<br>VD China C3<br>VD China C4<br>VD Viet Nam V1 |
|  |  | Thailand | 2003 | 1 | VD Viet Nam V1 |
|  |  | Viet Nam | 1996 | 2 | VD Viet Nam V1<br>VD Korea K1 |
|  |  | Myanmar | 2003 | 1 | VD Viet Nam V1 |
|  |  | Nepal | 2000 | 1 | **VD Nepal 2** |
| *V. destructor* | New host<br>*A. mellifera* | Japan | 1996 | 1 | VD Korea K1 |
|  |  | South Korea | 1996 | 3 | VD Korea K1 |
|  |  | China | 2001 | 5 | VD Korea K1 |
|  |  | Viet Nam | 1996, 1998 | 6 | VD Korea K1 |
|  |  | Myanmar | 2003 | 2 | VD Korea K1 |
| *V. jacobsoni* | Original host<br>*A. cerana* | India | 2002 | 2 | VJ Bengaluru 1 |
|  |  | Thailand | 2003 | 1 | VJ Laos L1 |
|  |  | Viet Nam | 2003-2004 | 5 | VJ Laos L1 |
|  |  | Malaysia | 1995 | 1 | VJ Malaysia 2 |
|  |  | Indonesia | 1998, 2002 | 6 | VJ Java 1<br>**VJ Lombok 2** |
|  |  | Papua New Guinea | 1997, 2008, 2015 | 4 | VJ PNG 1 |
| *V. jacobsoni* | Original host<br>*A. nigrocincta* | Indonesia | 1996 | 3 | VJ Java 1 |
| *V. jacobsoni* | New host<br>*A. mellifera* | Thailand | 1994 | 1 | **VJ North Thailand 4** |
|  |  | Papua New Guinea | 2008, 2014 | 9 | VJ PNG 1 |

Using the set of 2,091 SNPs detected across the mitogenome, we reconstructed phylogenetic relationships among *Varroa* lineages in each species (Figure 1). While exhibiting more genomic variability than previously reported, the mitogenome networks were consistent with the standard classification of *Varroa spp.* geographic lineages. We found that the consensus sequences of both species diverged 3.6%. Divergence within *V. destructor* samples was very low (0.5% with 85 SNPs), but moderate in *V. jacobsoni* with up to 1.3% (216 SNPs).

**Figure 1:**
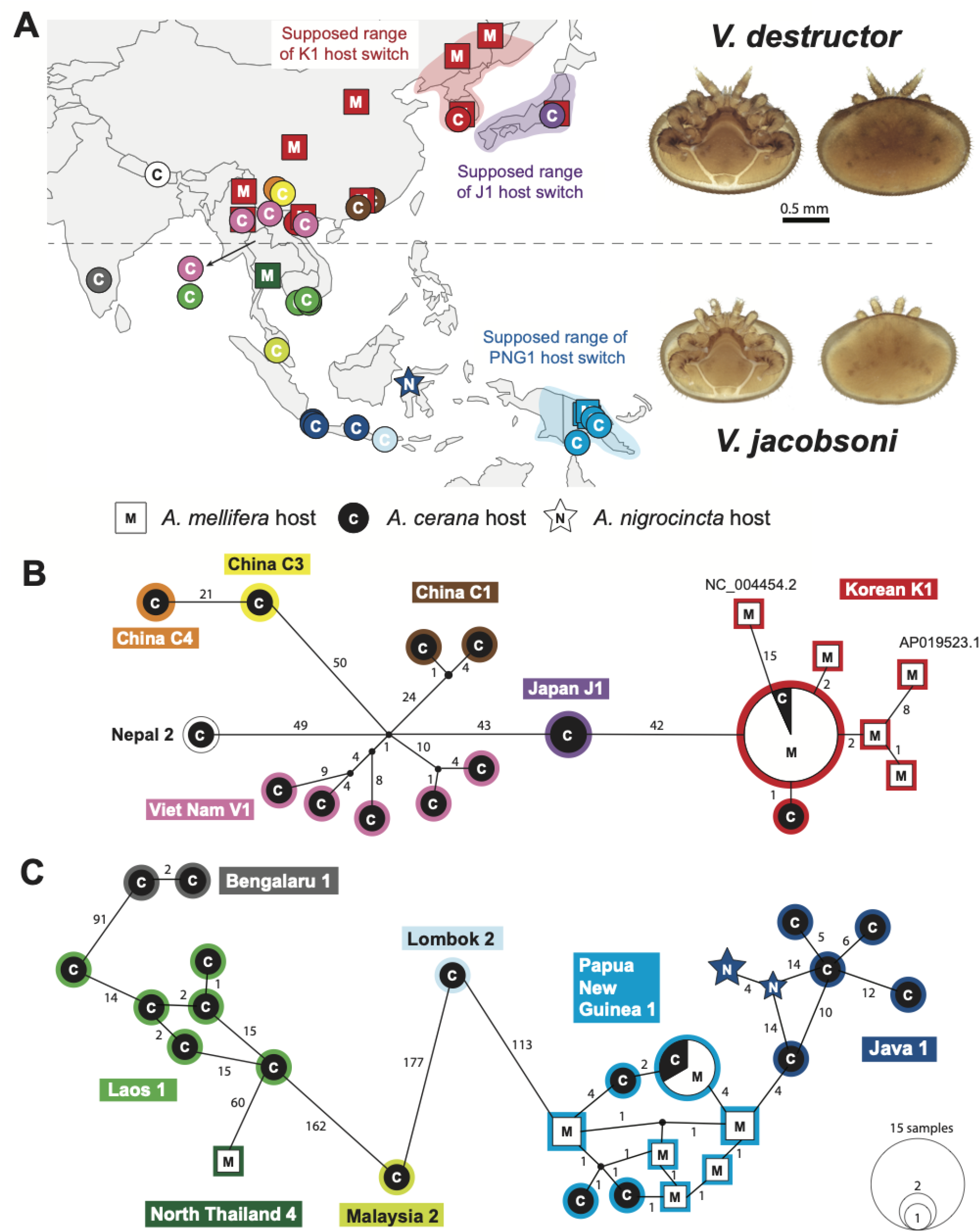
Mitogenomic phylogeographic networks uncover cryptic diversity in host-switched lineages. Both mite species exhibit geographic structure on their original host, with lineages classified using their *COX1* sequences (A). The color of each circle/square indicates a unique mitogenome sequence and size is proportionate to the sample size (B). Previous studies based on *COX1* found that *V. destructor* host-switched lineages largely belong to the so-called ‘Korean K1’ haplotype. However, the mitogenome-wide analysis revealed four six mitotypes on the new host, suggesting previously unreported switches by mites from the same geographic region (B). For the most recent jump in 2008, out of the seven lineages identified for *V. jacobsoni* only the known PNG1 succeeded in crossing the species barrier [4] (C). Yet, looking at the whole mitochondrial genome, it becomes evident that the host switch involved a number of independent female founders from closely related populations, or even the same population. Thus, mitogenomic data suggest a relatively diverse population of host-switched mites in both species.

We identified 18 distinct mitogenome sequences (hereafter named mitotypes) for *V. destructor* that clustered into seven lineages (Figure 1A and Table S2). Throughout our Asian sampling sites, all *V. destructor* mitotypes on *A. mellifera* belonged to the Korean K1 lineage, which was also found in one *A. cerana* sample (Figure 1B). We found three additional K1 mitotypes in China that were not previously reported on *A. mellifera* [20] (Table S1). We confirmed that the SNPs differentiating these K1 mitotypes were not artifacts of base calling by manually checking the read alignments. Interestingly, the two publicly available mitochondrial genomes of the K1 haplotype (France: NC_004454.2 and Japan: AP0195523.1) appear slightly divergent from the dominant K1 mitotype. Though we found the other reported host switched lineage (Japanese J1) in the source population from Japan, we did not detect it on *A. mellifera*. In addition, a re-analysis of 22 *COX1* sequences available on NCBI, combined with our new data also indicates diverse mitochondrial backgrounds of host-switched mites, both in *V. destructor* and *V. jacobsoni* (Figure S1).

Despite similar sample sizes for both species, we found a much higher genomic variability in the mtDNA in *V. jacobsoni* samples compared to *V. destructor*. Our analysis detected 27 unique mitotypes that belonged to seven lineages (Figure 1C and Table S2). Unexpectedly, we found that the misidentified mite on *A. mellifera* in Thailand (THV003_1) clustered with a new lineage, VJ North Thailand 4 (Table 1). This lineage was closely related to the North Thailand 1-3 *COX1* reference sequences (Figure S1) on and Laos 1 mitotypes (Figure 1C) [10,13]. All other *V. jacobsoni* mites collected on *A. mellifera* (*N* = 10), belonged to the previously reported host switched lineage Papua New Guinea 1. The median joining network showed six mitotypes found on A. *mellifera* were distinct from each other (up to 8 SNPs difference). We also found the two PNG 1 mitotypes detected during the host switch event (2008) that were retrieved several generations later in *A. mellifera* colonies (2014) (Table S1).

### Host-associated genomic differentiation following host shifts

We identified 2,728,471 biallelic SNPs in whole-genome sequences of *Varroa spp.* mites, after quality and coverage bias filtering. These variants revealed the loss of genetic diversity that accompanied the transition from the original to the novel bee host. For instance, a 2.8-fold reduction in polymorphic SNPs numbers was observed on the novel host *V. destructor* (N_VdesAcer_ = 1,219,990 vs N_VdesAmel_ = 429,420 SNPs) and 3.6 fold in *V. jacobsoni* (N_VdesAcer_ = 1,546,366 vs N_VdesAmel_ = 630,534 SNPs). Reduction of genetic diversity on the novel host was also observed in the average genomic nucleotide diversity π in both *V. destructor* (π_VdesAcer_ = 0.0020 vs π_VdesAmel_ = 0.0005) and *V. jacobsoni* (π_VjacAcer_ = 0.0014 vs π_VjacAmel_ = 0.0008). Genome scans using 20kb sliding windows showed that π values in the new host was overall lower than in the original host (Figure S2). Analysis of the genomic heterozygosity revealed extremely high levels of inbreeding coefficient in mite genomes with the lowest value F = 0.686 in *V. destructor* and F = 0.721 in *V. jacobsoni* (Table S3). However, when comparing *A. mellifera* and *A. cerana* host populations, we found no significant difference in F estimates for *V. destructor* (two-tailed t-test *p-value* = 0.263) and *V. jacobsoni* (two-tailed t-test *p-value* = 0.297). Such levels in both species were previously reported using microsatellites [4] and more likely resulting from incestuous *Varroa spp.* mating.

PCA further illustrated the loss of genetic diversity in host-switched mites coupled with genome-wide differentiation (Figure 2AC). Likewise, F_ST_ and D_xy_ pairwise estimates indicated strong differentiation between nuclear genomes from divergent lineages (Table S4). Within each species, high levels of genetic differentiation all along all seven chromosomes were detected between host-adapted lineages (Figure S2). Interestingly, when comparing the host-switched *V. destructor* specimens to native-host samples from the same mitotype (K1) there was far less genetic differentiation (F_ST_ = 0.004) compared with any other native-range population (F_ST_ > 0.6) (Table S4). Thus, host-switched populations still carry many of the genes present in the population of origin, a scenario consistent with rapid reproductive isolation post-switch. Similar patterns were obtained genome-wide for the larger sliding windows of 20kb of F_ST_ and D_xy_ estimates show that *A. cerana* and *A. mellifera* K1 mites diverged massively (Supplemental online markdown).

We found similar patterns of population structure and divergence among hosts and mtDNA lineages in *V. jacobsoni*, while host switch was more recent. In our study, we did not find evidence for quasi-clonality in either *V. jacobsoni* or *V. destructor* mites collected from *A. mellifera* populations.

### Ancestry of host switched populations and evidence of gene flow

NGSadmix analysis conducted on 1,276,602 unlinked biallelic SNPs confirmed the population structure observed between *A. mellifera* and *A. cerana* mites and helped to identify the source of host populations. At K = 2, population structure quickly emerged between *V. destructor* mites. J1 and K1 *A. cerana* mites were assigned to the same genetic cluster as all K1 *A. mellifera* mites (Figure 2B). Therefore, genome-wide sequencing confirmed that both K1 and J1 mite lineages contributed to the genetic diversity in host-switched populations. In *V. jacobsoni*, admixture estimates suggested that two large genetic clusters exist: Northern/Asian *V. jacobsoni* lineages differing from the South/Oceanian (Figure 2D). At K = 3, unreported spillover events were seen with the mite (THV003_1) found in *A. mellifera* in Thailand, which was genetically related to mites in Viet Nam (Laos 1). Here, population genomics supports the origin of *V. jacobsoni* host switch in Papua New Guinea to be exclusively from mites infesting *A. cerana* with PNG1 and Java 1 mitochondrial lineages.

**Figure 2:**
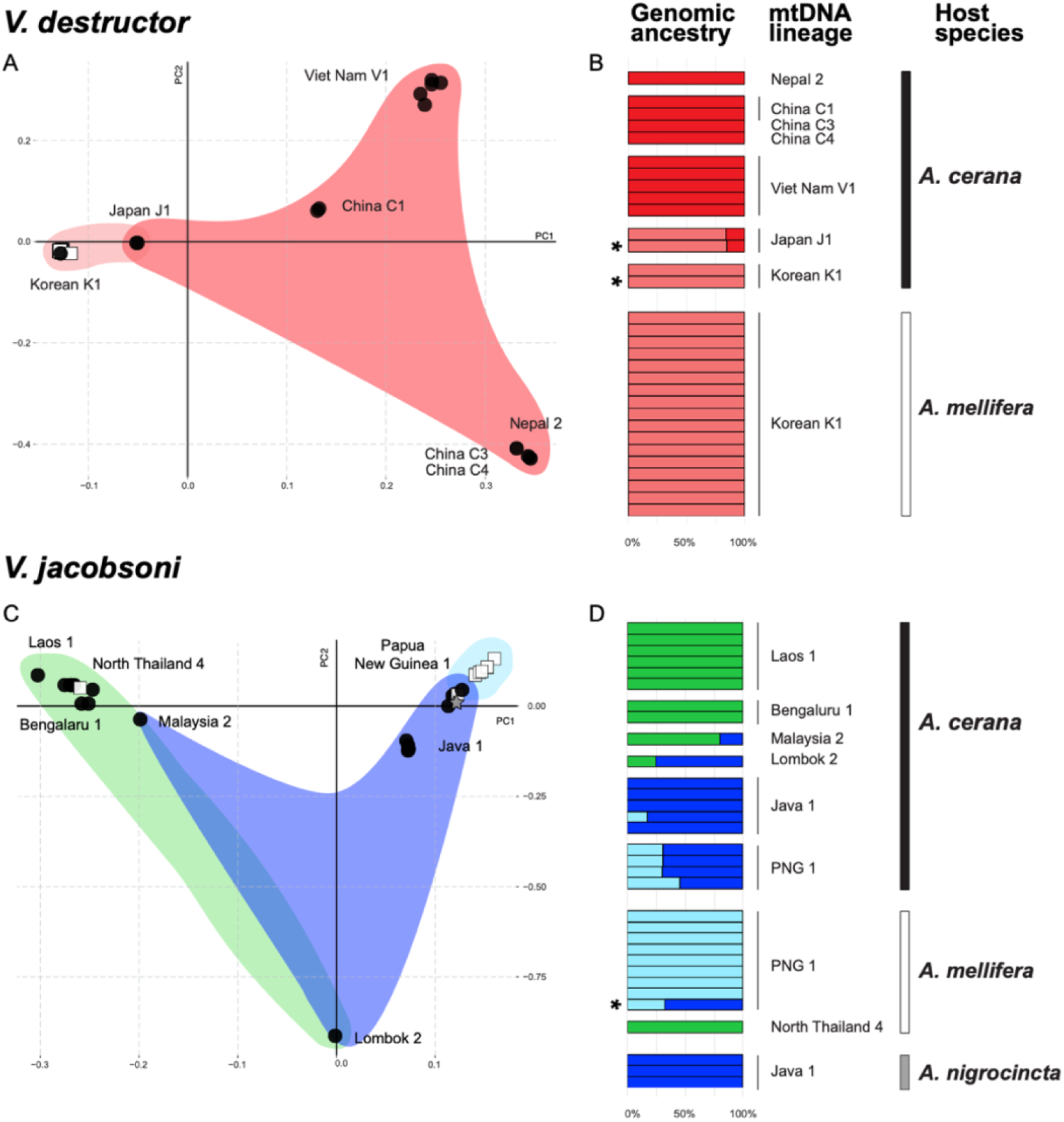
Loss of genetic diversity and rapid genomic differentiation occurred in spite of large founding size and migration between sympatric honey bee hosts. *V. destructor* populations on its original host *A. cerana* (black circle) show geographic structure across their native range (A). Envelopes around the PCA points are color-coded according to nuclear genetic ancestry (B). Host-switched populations (white square) are genetically homogenous throughout Asia and are most similar to the populations with the same mitotype (K1) (B). Despite a more recent host switch (∼2008), a similar pattern is observed for *V. jacobsoni* in South-Asia and Oceania, where the PNG1 lineage underwent a host switch (C). However, other switches appear possible elsewhere in *V. jacobsoni*’s range, such as the specimen collected from *A. mellifera* in North Thailand (green cluster) (D). In both species, there is evidence of mites drifting between hosts, as indicated by genetic analysis of their gut contents, indicated by asterisks. The overall picture for both species is similar, where reproductive differentiation after host switches is coupled with additional opportunities for gene flow via drifting, and potential for additional switches happening elsewhere in the range.

Recent migration and spillover events were confirmed by analyzing mite’s bee reads from inside the mite. We first evaluated this method with mite families (used for *de novo* mutation measurements) that were actively feeding at the time of collection and were able to accurately identify the host (Table S5). For other samples up to 0.18% of all reads from each mite library matched honey bee host mitochondrial reference genomes. Using this method we found that a *V. jacobsoni* mite collected on *A. mellifera* that did not cluster with the other host-switched mites based on nuclear DNA (PGV956_4) actually had predominantly *A. cerana* DNA inside her, suggesting a recent drifting event. While admixture estimates did not spot such outliers in *V. destructor*, the diet molecular analysis detected cases in Viet Nam (VNV475_1) and in Japan (JPV025_1) where mites from *A. mellifera* drifted back to *cerana*. Since *A. mellifera* and *A. cerana* honey bees present enough important morphological and behavioral differences to make confusing them impossible by experienced field workers, such parallel observations likely indicate that the mites migrated between hosts shortly before collection.

### Low *de novo* mutation rate from pedigree-based measurements

We examined whether the rapid parallel differentiation observed between *Varroa spp.* host populations was driven by high frequency of new mutations by estimating the *de novo* mutation rate, following a previously published approach [21]. As the *V. destructor* reproductive sequence cycle is well known (one haploid son, followed by up to four diploid daughters), we sequenced six diploid mothers and their respective sons at a coverage between 9x–47x for the mothers and 9x–54x for the sons. Before filtering, the number of detected mutations was 3,082 but there was extreme variation between samples in the number of potential mutations called. Notably mother-son pairs with less coverage contained several folds more mutation calls than mother-son pairs with higher coverage. For example, between Mom1 (47x) and Son1 (54x), 2 potential mutations were identified, whereas between Mom19 (9x) and Son19 (13x), 1582 potential mutations were identified. Samples with higher mutation calls also showed stronger evidence of clustering of mutation calls, which typically indicates false-positive mutation calls. For example, between Mom7 (9x) and Son7 (11x), 1454 potential mutations were identified; however, the null hypothesis that mutations were distributed to contigs in proportion to contig lengths was rejected (via a G-test) and the null hypothesis that mutations were distributed uniformly along the contigs was also rejected (via an Anderson-Darling test). Mom2-Son2 and Mom19-Son19 had similar results. After filtering to remove low quality mutation calls, two mutation calls remained, but showed low coverage and were discarded as unreliable.

We conducted a set of simulations to deal with coverage bias issues in the data set. Out of 6,000 simulated mutations, 3,464 could be detected by our pipeline, indicating that our recall rate was approximately 58%. The total haploid length of the 7 chromosomes considered in the mutation scan was 362,833,908 base pairs. Given that we found no mutations among 6 mother-son pairs, we estimate that the mutation rate in *V. destructor* is less than 8 × 10^−10^ per bp per generation 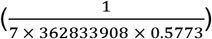 or less than 0.28 mutations per son per generation.

### Estimating of host switch demographic history and founding population size using nuclear DNA

We aimed to further parameterize the demography of host switches detected in mitochondrial data using coalescent-based modeling based on nuclear DNA. Demographic inference using fastsimcoal2 was conducted using an isolation with migration scenario (Figure S3). For *V. destructor*, 32 haploid mite genomes were sequenced from *A. mellifera* host and 22 haploid mite genomes were sequenced from *A. cerana* host. For *V. jacobsoni*, 18 haploid genomes of *A. mellifera* mites and 6 haploid genomes of *A. cerana* mites from Papua New Guineas were projected (excluding the older sample from 1997). Given that we were only able to estimate the upper limit of *de novo* mutation rate, we explored whether lower mutation rate improve the observed maximum likelihood distribution with 100 replicate simulations (Figure S4). We examined rates as low as 1.0 × 10^−11^, at the lower limit of rates in arthropods [22]. We also tested whether including an inbreeding coefficient would help improving the model likelihood by adding 70% of inbreeding (minimum detected in both species). However, lower mutation rates did not improve model fit for both *V. destructor* (Kruskal-Wallis chi-squared = 497.61, df = 494, *p-value* = 0.446) and *V. jacobsoni* (Kruskal-Wallis chi-squared = 499, df = 498, *p-value* = 0.479) (Figure S4). The inbreeding coefficient did also not significantly improve the likelihood with equal mutation rate (available in online supplemental markdown). In addition, 2D-joint SFS visual inspection did not reveal striking differences, and the parameter estimates overall were qualitatively similar over the range of mutation and inbreeding parameters we examined.

Hence, demographic parameters and their bootstraps values were estimated using μ = 8 × 10^−10^ per bp per generation and no inbreeding value. Mean values and 95% confidence intervals for each species are graphically summarized in Figure 3. The older host switch with *V. destructor* was estimated to have occurred 887 generations before sampling (∼88 years ago assuming 10 generations / year [23]) with a founder effect lasting for 177 generations. Estimates suggested that 156 [CI95 = 146-165] haploid genomes of *V. destructor* contributed to the founding event in Asia. From that point on, the novel mite population grew quickly on *A. mellifera* and diverged from the sympatric *A. cerana* source populations. Our results suggest low but continuous gene flow between hosts. Despite being a more recent event, estimated around 183 generations ago (∼18 years ago), demographic parameters for *V. jacobsoni* also supported a large founding size reaching 138 haploid mites. The effective population size of *V. jacobsoni* mites on *A. mellifera* was lower than the source population, in contrast to estimates for *V. destructor*.

**Figure 3:**
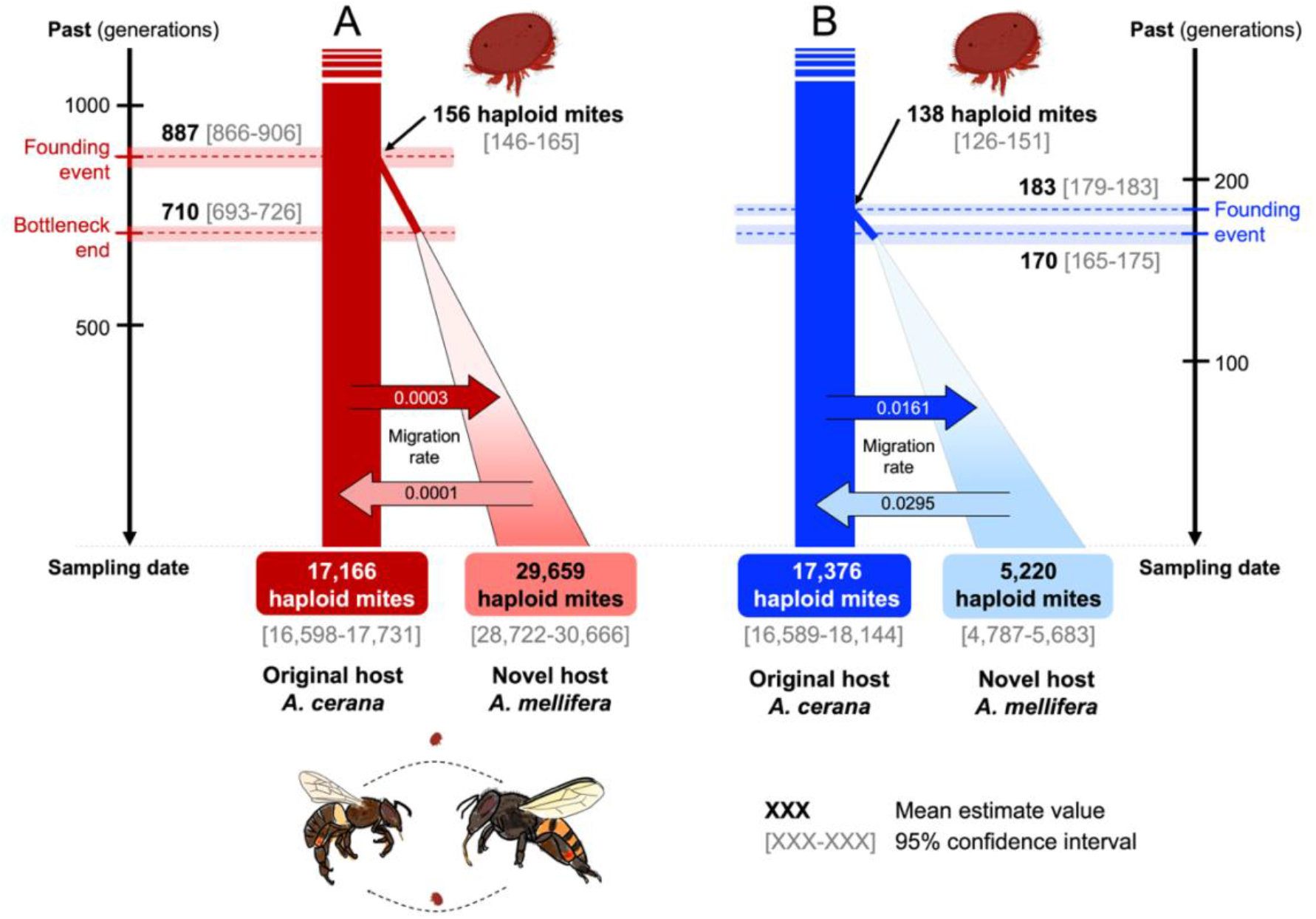
Multiple jumps to *A. mellifera* and isolation with migration likely allowed *V. destructor* and *V. jacobsoni* to successfully switch hosts and persist. Graphical illustration of the inferred scenario with mean parameter estimates in bold and associated 95% confidence interval in [light gray] with isolation with bidirectional migration (A) for *V. destructor* and (B) for *V. jacobsoni*. Estimated population sizes are given in haploid genome size. Migration rates were kept as default values in haploid genome per generation. The coalescent model reconstructed timings of expansion that are consistent with known observations. Yet it suggests that the host switch was not instantaneous and may have involved a period of progressive adaptation before the mites spread beyond their source populations.

## Discussion

Virtually every living species has at least one parasite, making parasitism perhaps the most successful mode of life. Most studies examine parasitism from either a macroevolutionary perspective, for instance, through host-parasite co-phylogenies, or at the population level, by examining patterns of differentiation between parasites and hosts [24,25]. Theoretical modelling links the two scales, suggesting that specialization coupled with trade-offs in performance on alternative hosts should lead to speciation [26]. Yet, empirical observations of this process remain scarce, particularly of host switch demography, which includes key parameters such as population sizes during switches that are necessary to ensure sufficient evolutionary potential for parasites [27]. For this reason, pandemic spread of *Varroa spp.* among honey bees has long been puzzling, given their previously assumed “quasi-clonal” structure in the invasive portion of their range [9]. Using high-resolution mitochondrial and nuclear genomic data from sympatric populations, we show that in both *V. destructor* and *V. jacobsoni,* (a) genetic bottlenecks were far less severe than previously estimated and (b) while gene flow was greatly reduced post-host switch, consistent with models of incipient speciation that may accompany acquisition of novel hosts, gene flow has not completely stopped. Our data highlight the importance of genetic diversity during initial stages of parasitic host switches.

While loss of genetic diversity is common in parasites, it often occurs in the parasite’s introduced geographic range [28,29]. Consequently, parasites may possess sufficient genetic diversity to parasitize their hosts. For example, the fungal parasite responsible for ash dieback in Europe arrived from Asia, and while bottlenecked to only two haplotypes, it nonetheless maintains adaptive diversity in key host interaction genes [14]. To the best of our knowledge, our study is unique in quantifying demographics within the native range of the parasite, where it has opportunities for additional gene flow from genetically diverse sympatric populations. At least for *Varroa spp.,* the initial host switch requires an unexpected amount of genetic diversity at the point of the switch, but rapidly leads to reproductive isolation between sympatric populations on novel and original hosts. The amount of differentiation appears to increase over time, being greatest in *V. destructor*, which switched ∼50 years earlier than *V. jacobsoni*. *Varroa spp.* coevolve quickly with their new hosts [30,31] and such sympatric isolation could ultimately lead to speciation.

Post host-switch speciation can occur in the presence of gene flow, if specialization is adaptively favored and selection acts on many genomic regions [32,33]. By promoting genomic heterogeneity and introducing beneficial mutations [34], gene flow can ameliorate lost diversity due to selection or inbreeding [35,36], both of which would impact host-switched *Varroa spp.* populations. Alternatively, *de novo* mutation rates could contribute to population genetic variation, but our direct estimation of mutation rate did not show exceptionally high mutation rates (< 8 × 10^−10^ per bp per generation), compared to other arthropods [22,37]. Consequently, immigration may be the primary means of introducing new alleles into populations on the novel host during early stages of the host switch. Recent work has shown experimentally that gene flow between can occur between *V. destructor* populations on the two hosts in the native range [20]. This observation, together with migration between host species detected in this study provides a mechanism by which gene flow occurs – through occasional co-infestations by females from different populations. In this case, even if a female is not fully adapted to a host, she may still be able to propagate her genes by laying a male even without mating [38], who can then mate with the other female’s daughters [39,40].

While host switch events by specialized parasites were once assumed to be rare, this assumption is increasingly challenged [41,42]. Ecological fitting theory suggests that shifts could readily occur in species with a pre-existing ability to use novel hosts [43]. Host switches in *Varroa spp.* appear to fit this model. While the western honey bee has been introduced throughout the *Varroa spp.* native range, mitochondrial data, which are geographically informative (Figure 1), suggest that switches occurred only in Korea, Japan, Philippines (*V. destructor*), Papua New Guinea and possibly Thailand (*V. jacobsoni*) [4,11–13]. *A. cerana* subspecies are strongly differentiated geographically [44–49], and *Varroa spp.* mitotypes mirror host biogeography and subspecies distribution [10,12,13,50]. This suggests that *Varroa spp.* populations may vary in traits, such as host specificity, as a result of their previous coevolutionary interaction with local *A. cerana* subspecies. This may have allowed some populations to switch, while others were unable to do so, in spite of available novel hosts [51]. However, the dynamic nature of the interaction does not preclude additional switches in the future.

One potential limitation of our data set is its sparse geographic and temporal sampling, given the size of the native range. While small sample size is enough to retrace ancestral events, large sample sizes can increase the confidence in model selection and parameter estimates for recent demographic events [52]. Yet, past scenario and demographic estimates were consistent with mite biology and reflected direct observations made during their invasion of *A. mellifera*. Furthermore, while our data set captured most of the described *Varroa spp.* mitochondrial lineages and even discovered new ones [3,9,13,53], some reported mitotypes were missing. In the future, incorporating larger sample size from modern source populations could help better estimate each lineage genetic contribution to host switch success. In addition it would allow to estimate how host switch may have affected native diversity as for the case of J1 lineage displacement [54,55]. Despite these limitations, our analysis correctly reconstructed known aspects of *Varroa spp.* host switch demography, such as the times of the switches. However, these times need to be regarded as approximate, since generation times may fluctuate due to a variety of factors, such as seasonality or brood availability. We relied on a previous approximation of 10 generations per year [23]. Finally, the use of historical samples, which were collected over the span of approximately a decade, may also affect parameter estimates in some way. As a result, we caution against over-relying on the apparent precision of the numerical estimates (Table 2). Nonetheless, data from both mitogenome sequencing and coalescent analysis all point to a relatively large founding population size of the *Varroa spp.* mites, the existence of migration and striking parallels in the demographics of host switches between the two species (Figures 1 and 3).

**Table 2:** Parameter estimates for the most likely demographic scenarios with isolation with migration and population expansion for each species jump. As stated in fastsimcoal2 manual, these upper-bound priors indicated with an asterisk were not bounded by the analysis.

|  | Priors |  | *V. destructor* |  | *V. jacobsoni* |  |
| --- | --- | --- | --- | --- | --- | --- |
|  | Distribution | Range | Mean value | CI 95% | Mean value | CI 95% |
| **N<sub>VAC1</sub>**<br>(haploid genomes)<br>At present, mite effective population size on original host *A. cerana* | uniform | [2; 20,000*] | 17,166 | [16,598; 17,731] | 17,376 | [16589; 18144] |
| **N<sub>VAM0</sub>**<br>(haploid genomes)<br>At present, mite effective population size on novel host *A. mellifera* | uniform | [2; 20,000*] | 29,659 | [28,722; 30,666] | 5,220 | [4,787; 5,683] |
| **N<sub>BOTAM</sub>**<br>(haploid genomes)<br>Past mite effective founding population size during host switch | uniform | [2; 1,000*] | 156 | [146; 165] | 138 | [126; 151] |
| **T<sub>JUMP</sub>**<br>(generation)<br>Time of new founding population split by host switching | uniform and bounded | [1; 1,000] | 887 | [866; 906] | 183 | [179; 187] |
| **T<sub>BOTEND</sub>**<br>(generation)<br>End of the founder effect period | uniform and bounded | [1; 1,000] | 710 | [693; 726] | 170 | [165; 175] |
| **G<sub>AM</sub>**<br>(haploid genomes per generation)<br>Growth rate of the expanding newly founded mite population (from present to past) | uniform | [-1; 0*] | -0.0014 | [-0.0015; -0.0013] | -0.4636 | [-0.4895; -0.4399] |
| **M<sub>ACToAM</sub>**<br>(haploid genomes per generation)<br>Migration rate of parasite spillover (original toward new host) | log uniform | [1x10<sup>-6</sup>; 100*] | 0.0003 | [0.0003; 0.0003] | 0.0161 | [0.0154; 0.0168] |
| **M<sub>AMtoAC</sub>**<br>(haploid genomes per generation)<br>Migration rate of parasite spillback (new toward original host) | log uniform | [1x10<sup>-6</sup>; 100*] | 0.0001 | [0.0001; 0.0001] | 0.0295 | [0.0283; 0.0309] |
| **Mutation rate (fixed-parameter)** | - | 8.0 x 10<sup>-10</sup> |  |  |  |  |
| **Inbreeding (fixed-parameter)** | - | none |  |  |  |  |

Our findings highlight the dynamic and ongoing nature of host switches in the native range, and the need to better understand native mite populations. Mitochondrial DNA has already detected the likely presence of several host switches, such as those in Philippines in 2015 [12] and the presence of new haplogroups in Eastern Europe [56], indicating that populations of *Varroa spp.* are continuously testing *A. mellifera* as a new host, and may have spread without our awareness. Additionally, while this work provides insight into the initial host switch, the demographics of the subsequent worldwide pandemic remain largely unknown, for instance the critical population size necessary to establish a regional infestation. Future research on global demographic parameters such as genetic diversity and gene flow is crucial [57]. Since *V. jacobsoni* presents striking parallels to *V. destructor*, this knowledge could be applied to forecast and prevent its spread.

## Materials and Methods

### Mite sampling on original and novel honey bee hosts

We sampled and sequenced mites for two purposes: 63 *Varroa spp.* females were collected for population genomics (described in this sub-section) and 12 specimens were used for *de novo* mutation rate estimation (described in a later sub-section). We sequenced adult females throughout the *Varroa spp.* native ranges where original and novel hosts occur in sympatry (for population genomics: *V. destructor N* = 31 and *V. jacobsoni N* = 32; Table 1). For *V. destructor,* specimens were collected between 1996 and 2003 as part of the taxonomic revision and diversity survey carried by Anderson and Trueman [3] and Navajas *et* al. [11] and included 14 samples from *A. cerana* and 17 from *A. mellifera* collected from Japan, South Korea, China, Thailand, Viet Nam, Myanmar and Nepal (see interactive maps https://maevatecher.github.io/varroa-host-jump/#sampling-distribution). For *V. jacobsoni*, female specimens were obtained from Anderson and Trueman [3] (1994-2004) and Roberts *et al*. [4] (2008-2015) surveys of host shifts reports in Papua New Guinea. A total of 19 adult female *V. jacobsoni* mites were collected from *A. cerana* across different areas in India, Thailand, Viet Nam, Malaysia, Indonesia and Papua New Guinea. In contrast to *V. destructor*, *V. jacobsoni* infestation on *A. mellifera* is still restricted to Papua New Guinea from which 10 samples were collected. Additionally, we included three *V. jacobsoni* mites collected from an alternative original host *A. nigrocincta* found in Indonesia. *A. nigrocincta* is not believed to be the source of the host-switched mites, so these samples were included as an outgroup.

Sampling complete details regarding host, location and year are available in Table S1. Exact geographical coordinates were not always available for samples collected before 2008, and were approximated by the locality provided, or published survey maps [3,11,53,58]. All individuals were mature sclerotized females collected from single colonies and were preserved in individual Eppendorf tubes. Each collection tube was stored with 70% ethanol and kept at −20°C at the CSIRO in Canberra, Australia.

### DNA extraction and whole genome resequencing

Mites were surface sterilized by cleaning them in absolute ethanol using a sterile brush to remove any external debris, and then gently shaken in a 2.0 mL Eppendorf filled with absolute ethanol. Each mite was then dried for 10 sec on a sterile paper towel before being placed in a 1.5 mL Eppendorf tube in liquid nitrogen. Genomic DNA was extracted from each mite by crushing the whole body, using a sterile pestle to obtain a fine powder and processed with a QIAamp DNA Micro Kit (© Qiagen) following the manufacturer’s instructions. Final elution volume was 15 μL. Total dsDNA was measured using a Qubit™ 4 Fluorometer with an Invitrogen dsDNA HS Assay Kit.

For population genomic samples, short-inserts of 150-bp paired-end libraries were prepared for each individual using a Nextera XT DNA Library Preparation Kit (Illumina ®). Size-selection and cleanup were accomplished using CA-magnetic beads (Dynabeads® MyOne Carboxylic Acid, Invitrogen), and 11-11.5% PEG 6000 (Sigma-Aldrich © LLC). Library quality and size were assessed using a Bioanalyzer High Sensitivity DNA kit (Agilent). Libraries were run on HiSeq 4000 and NovaSeq6000 in 150 bp × 2 paired-end mode (Illumina ®) at the OIST Sequencing Center. Biosamples and DRA accession as well as sequencing coverage for each sample are provided in Table S1.

### Host DNA contents in mite whole-body metagenomics

We cross-checked honey bee host identities reported during sampling with host read identity to detect potentially migrating mites. Mites feed on honey bees during the phoretic phase [59] and maintain a consistent feeding regimen by consuming ∼1 μL of host fluid (digested fat-body and haemolymph) per day [60]. Therefore, we assumed that host DNA would be retrieved from crushed whole-mite tissues. Mitochondrial DNA was targeted, as it is more abundant than nuclear DNA. We mapped raw fastq reads on honey bee host mitochondrial reference genomes using [NC_001566.1] for *A. mellifera ligustica* [61] and [NC_014295] for *A. cerana* [62]. The number of reads mapped to either one of these honey bee host reference genomes was counted and compared to sampled host identities.

### Data filtering, mapping and genotype calling

Commands used for each analysis step are available on our Snakemake script [63] available on https://github.com/MaevaTecher/varroa-host-jump. Briefly, we assessed demultiplexed fastq read quality using FastQC [64]. We then mapped reads to the *V. destructor* reference genome on NCBI [GCF_002443255.1] [65] separately from the complete mitogenome [NC_004454.2] [66] using the soft-clipping and very sensitive mode of NextGenMap v0.5.0 [67] (following a comparison with Bowtie2 v2.6 [68]). Reads were sorted and duplicates were removed using SAMtools [69], and subsampled to a maximum coverage of 200 using VariantBam [70] to speed up processing. Mapping rates and reads depths were computed from the generated BAM files.

We generated three data sets for the mapped reads depending on analysis requirements, following Yamasaki *et al.* [71]. First, we obtained a “*SNP-only dataset*” containing only variant sites using FreeBayes v1.1.0 [72] with the following parameters: minimum mapping = 10, minimum base quality = 5, use of the four best SNP alleles, and no populations prior. In order to correct for coverage bias and sequencing artefacts in problematic regions, we estimated twice the mean reads depth along the seven main chromosomes. Subsequently, variants were filtered using using VCFtools v0.1.12b [73]. The “*SNP-only dataset*” resulted in 2,728,471 SNPs. Second, we computed an “*all-sites datase*t” by calling both variants and invariants sites using BCFtools v1.9 mpileup [74]. We removed indels and sites at over twice the computed mean depth, as well as sites with any missing data. We excluded sites that were not placed on the seven chromosomes. After filtering, the “*all-sites dataset*” resulted in 2,130,335 SNPs and 120,279,163 monomorphic sites. Third, we obtained a “*mtDNA SNP-only dataset*” by calling polymorphic sites only on the mitochondrial genome NC_004454.2. We used FreeBayes v1.1.0 with strict quality parameters. The dataset resulted in 2,091 SNPs. These were validated in the *COX1* region by Sanger sequencing (see next section).

### Comparative mitogenomes analysis and lineages identification

We used the “*mtDNA SNP-only dataset*” for mitochondrial variation genomic analysis by converting and generating individual nucleotide sequences using the option *vcf2fasta* in vcflib scripts (github.com/vcflib). Nucleotide sequences were screened and aligned (16,476 bp) using Geneious Prime® 2019.2.3 together with the reference mitochondrial sequences NC_004454.2 (*V. destructor* on *A. mellifera*, France [66]) and also, to avoid problems with reference bias, from AP019523.1 (*V. destructor* on *A. mellifera*, Japan [75]). Mitotype diversity for each *Varroa* species and distribution in hosts was assessed using the median joining network method from SplitsTree5 [76]. Divergence levels between *Varroa* species was estimated by comparing the whole-mitochondrial consensus sequences with Geneious Prime®.

Standard methods exists to determine *Varroa* species and lineage (also named haplogroup) identity by using the following mitochondrial markers: 1) a 458-bp fragment from the *COX1* gene [3,77] and 2) a concatenated 2,696-bp fragment including *COX1*, *ATP6*, *COX3*, and *CYTB* [11]. Thus, we determined the lineage (e.g. Korean K1, Japan J1, *etc.*) of each mitotype by extracting the *COX1* barcoding region of interest and aligning it together with 60 unique *COX1* reference sequences from NCBI using ClustalW and manual check. A neighbor-joining tree was computed on the fasta alignment using IQ-tree [78] and exported with iTOL [79]. Following *Varroa* revised nomenclature [5], a sequence was considered from a known lineage only if 100% identical to reference *COX1* haplotype. The same process was used for sub-lineages (e.g. Korean K1-1/K1-2, Korean K1-3,…) identification by with the *COX1-ATP6-COX3-CYTB* concatenated region sequences which were aligned with 22 available reference concatenated sequences for *V. destructor* only [11,56,80].

To ensure that filtering of the “*mtDNA SNP-only dataset*” did not remove existing and known variants, we also sequenced the *COX1* gene using Sanger sequencing using primers 10kbCOIF1 and 6,5KbCOIR [11]. PCR reactions were carried out in 25μL containing 5 μL of 5X Phusion® HF buffer; 0.5 μL of dNTP mix (10mM); 0.25 μL of Phusion® High-Fidelity DNA Polymerase (NEB); 1.25 μL of each oligo primer 10kbCOIF1and 6,5KbCOIR (10mM); 1 μL of template DNA (0.5 ng/μL) and Milli-Q water. Samples were denatured at 98°C for 30 sec, and then PCR was performed for 35 cycles of 10 sec denaturation at 98°C, 15 sec of annealing at 59°C and 15 sec of extension at 72°C with a 5 min final elongation at 72°C. DNA amplification success was visualized by loading 3 μL of PCR product with 3 μL of loading dye on 1% agarose gel (110V for 20 min). PCR products were then cleaned-up using Dynabeads® MyOne Carboxylic Acid, CA-beads (Invitrogen) and 19% Polyethylene glycol PEG. Directly, fragments were sequenced in the two directions using the BigDye™ Terminator v3.1 Cycle Sequencing Kit (Thermo Fisher) in a capillary sequencer Applied Biosystems 3730xl DNA Analyzer (Thermo Fisher). FASTA sequences generated by mitogenome mapping and Sanger sequencing were then aligned and checked for differences.

### Genome-wide variation and analysis of divergence

We examined levels of genetic diversity and differentiation between host-adapted population of each *Varroa* species. To investigate differentiation among populations, we first measured the absolute divergence D_xy_ which requires both variants and invariants sites [81]. We also investigated levels of standing genetic variation (π). Genome scans were computed using a sliding window method on the “*all-sites datase*t” (50kb window, sliding every 20kb and containing a minimum of 100 SNPs). This method was implemented using the *parseVCF.py* and *popgenWindow.py* python scripts (github.com/simonhmartin/genomics_general) [82].

Using the biallelic “*SNP-only dataset*”, we also estimated the genome-wide differentiation by calculating Weir and Cockerham’s F_ST_ per site using VCFtools. Subsequently, we assessed the genetic structure within species using a principal component analysis (PCA) performed with the R package vcfR [83]. To further investigate the population structure and related ancestry, we conducted an incremental K step analysis using NGSadmix [84]. We reduce the effect of linkage disequilibrium between SNPs by conducting a pruning using PLINK v1.90b3 [85] on the “S*NP-only dataset*” with the following parameters: --indep-pairwise 20 10 0.5. After removing linked SNPs, the pruned dataset containing 1,276,602 SNPs was converted into BEAGLE format and directly used in NGSadmix. A total of 10 replicates per K, from 2 to 20, were run and the model with the highest likelihood was selected for plotting each K. The number of genetic clusters among and within species was determined following guidelines in [86,87] and making biological sense.

### Mutation rate estimation from mother-son pairs

Coalescent analysis requires an estimation of mutation rate, yet none is available for closely related species. To do this we collected and sequenced six *V. destructor* diploid mothers and their haploid sons for this purpose. These samples were collected from individual capped cells containing *A. mellifera* red-eye drone pupae. Each family was selected under the following conditions: 1) each cell was only infected by a single mother-foundress mite, 2) a *V. destructor* family at this stage should contained three to four offspring, including one haploid son and two to three diploid daughters, 3) the son was at the deutonymph or adult phase to avoid misidentification with protonymph sisters. We collected three families and preserved in absolute ethanol in February 2018 (#1, #2, and #7), and three more in May 2018 (#15, #17, and #19) from the same colony at the OIST Ecology and Evolution experimental apiary.

Libraries were prepared using an NEBNext® Ultra™ II FS DNA Library Prep Kit (New England Biolabs, Inc) with a fragmentation size incubation time of 15 min at 37°C, and three PCR cycles. Libraries were cleaned up using CA-magnetic beads (Dynabeads® MyOne Carboxylic Acid, Invitrogen), and 17% PEG 6000 (Sigma-Aldrich © LLC). Libraries were pooled and sequenced on two lanes of HiSeq 4000 (Illumina ®) at the OIST sequencing center.

Mutation rate was estimated from GATK variant calls [88] using DeNovoGear’s dng-call algorithm [21] which can model *Varroa’*s haplodiploidy sex-determination system. Single-nucleotide mutations were called on the 7 largest contigs in the Vdes_3.0 genome, avoiding sites that were within 100 bp of an indel. Mutation calls were de-duplicated, retaining only sites that were biallelic and not part of a mutation cluster (with 100 bp of another call or part of a long-run of calls from the same sample). After deduplication, the remaining calls were filtered such that P (denovo| data) was high (DNP >= 0.75) and the fit to the model was good (LLS >= −3). DNP (de-novo probability) and LLS (log-likelihood scaled) are both per-site statistics generated by dng-call. The denominator for the mutation rate analysis was estimated by simulating mutations at 1,000 locations in each son and calculating what fraction of these simulated mutations was recovered by our pipeline. A VCF was generated for these locations, and mutations were simulated by changing the son’s haplotype to one of the other three bases. ALT and AD fields were updated as needed.

### Demographic analyses of host switch using SFS

We inferred the demographic history of both *V. destructor* and *V. jacobsoni* mites with the coalescent simulator fastsimcoal v2.6 using the site frequency spectrum (SFS) [89]. Demographic inferences were computed independently for each species and samples were subsetted following genome-wide analysis results while considering geographical sympatry and continuity. For *V. destructor* (*N* = 27), we excluded Nepal and the Japanese mites (continental island) whereas for *V. jacobsoni* (*N* = 12), we included only mites from Papua New Guinea (host switch region). To reduce the effect of selection which can bias the SFS <u>(71)</u>, we kept SNPs and invariants from the “*all-sites datase*t” that were at least 50 kb away from any annotated genes regions (VCFtools exclude-bed option). The 2D-joint folded-SFS was computed for each species/host population on the filtered 12,594,802 sites including 224,568 SNPs, using the *vcf2sfs* R scripts (github.com/shenglin-liu/vcf2sfs) [90].

We considered a scenario consistent with the known history of *Varroa spp.* jumps (Figure S3). We incorporated the following demographic parameters (Table 2): the estimated effective population size for *Varroa* mites parasitizing *A. cerana* N_VAC1_ (in haploid genomes) to be stable before, during, and after the host switch. On the other hand, the mite population size on the new host *A. mellifer*a expanded from N_BOTAM_ or not with a growth rate G_AM_ after the host switch to reach modern population size N_VAM0_ (in haploid genomes). The host switch founder event occurred at a time T_JUMP_ (in generations) and ended at a time T_BOTEND_ (in generations). Finally, in a case of bidirectional migrations due to sympatry we estimated M_AMtoAC_/M_ACtoAM_, to be the proportion of haploid genomes migrating from one population to another.

We ran 100 replicates using the observed SFS as follow: a minimum of 20 loops (--minnumloops 20) and a maximum of 150 loops (e.g., ECM cycles, --numloops 100) were performed to estimate the parameters, with one million coalescent simulations per loop (--numsims 1,000,000), and a 0.001 minimum relative difference in parameter values estimated by the maximum composite likelihood (--maxlhood 0.001). The replicate with the highest likelihood set of estimated parameters was retained for model comparison.

To ensure that the upper limit genome-wide mutation rate µ = 8.0 × 10^−10^ previously used was adequate to estimate parameters from the observed SFS, we tested different µ levels. For this we ran scenario 4 in the same conditions as for scenario choice but µ ranging from 8.0 × 10^−10^ to 1.0 × 10^−11^. Additionally, fastsimcoal2.6 offers the possibility to input inbreeding coefficient. As *Varroa spp.* populations showed high inbreeding, we also ran scenario 4 with (*F* = 0.7, preliminary tests also ran with 0.8 and 0.9) and without inbreeding coefficient. For all these conditions combined, we ran 100 replicates for each of the 10 combinations per species.

Finally, the overall best estimate parameters set was used to generate 100 pseudo-observed SFS (for a similar number of polymorphic SNPs) for parametric bootstraps. We repeated the parameters estimation for each pseudo-observed SFS and kept the best run. The top 100 runs estimated parameters values were used to calculate 95% confidence intervals. The mean and 95 percentile confidence intervals were computed using the ‘boot’ R package [91]. Finally, the fit of the best expected SFS was visualized against the observed using SFStools R scripts (github.com/marqueda/SFS-scripts).

## Data Availability

All sequencing data generated for this study have been deposited in the DNA Databank of Japan and transferred to NCBI under the bioproject PRJDB9195 (https://www.ncbi.nlm.nih.gov/bioproject/650181) with the accession series DRR209082-DRR209125 and DRR212369-DRR212380. All customs scripts and pipeline developed using Snakemake, Rmarkdown have been made available on GitHub: https://github.com/MaevaTecher/varroa-host-jump. Demographic inferences input files and FASTA sequences alignment (mtDNA) are made available in the same repository. Variant calling files, genome indexing files and input lists are readily available from the following DRYAD repository: https://doi.org/10.5061/dryad.mgqnk98x1.

## Author Contributions

MAT and ASM designed research, analyzed population genetics data and wrote the manuscript. MAT processed the samples in the wet lab until library preparation and ran the demographic inferences. JMKR collected and provided samples from the mite CSIRO collection and participated in the data interpretation. RAC analyzed data, wrote the manuscript, provided reproducible online resources and data for the estimation of mutation rate.

## Acknowledgments

MAT’s research was supported by a postdoctoral fellowship from the Japan Society for Promotion of Science (JSPS) (P19723), Kakenhi Grant-in-Aid, (19F19723) and the Okinawa Institute of Science and Technology (OIST). ASM was supported by a Future Fellowship from the Australian Research Council (FT160100178) and a Kakenhi Grant-in-Aid for Scientific Research from the JSPS (18H02216). JMKR was supported by the Australian Centre for International Agricultural Research (ACIAR) and the Australian Department of Agriculture, Water and Environment (DAWE). RAC was supported by the National Institutes of Health (R01-HG007178). We are grateful to Jo Si Lay Tan and Lijun Qiu for their advice and guidance in developing the wet lab workflow for *Varroa* mite sequencing. We also would like to thank the OIST Sequencing Center for assisting us in the sample sequencing. We wish to thank Steven D. Aird, technical editor, for reviewing and improving our manuscript.

## Ethics approval and consent to participate

Not applicable.

## Consent for publication

Not applicable.

---

**Figure S1:**
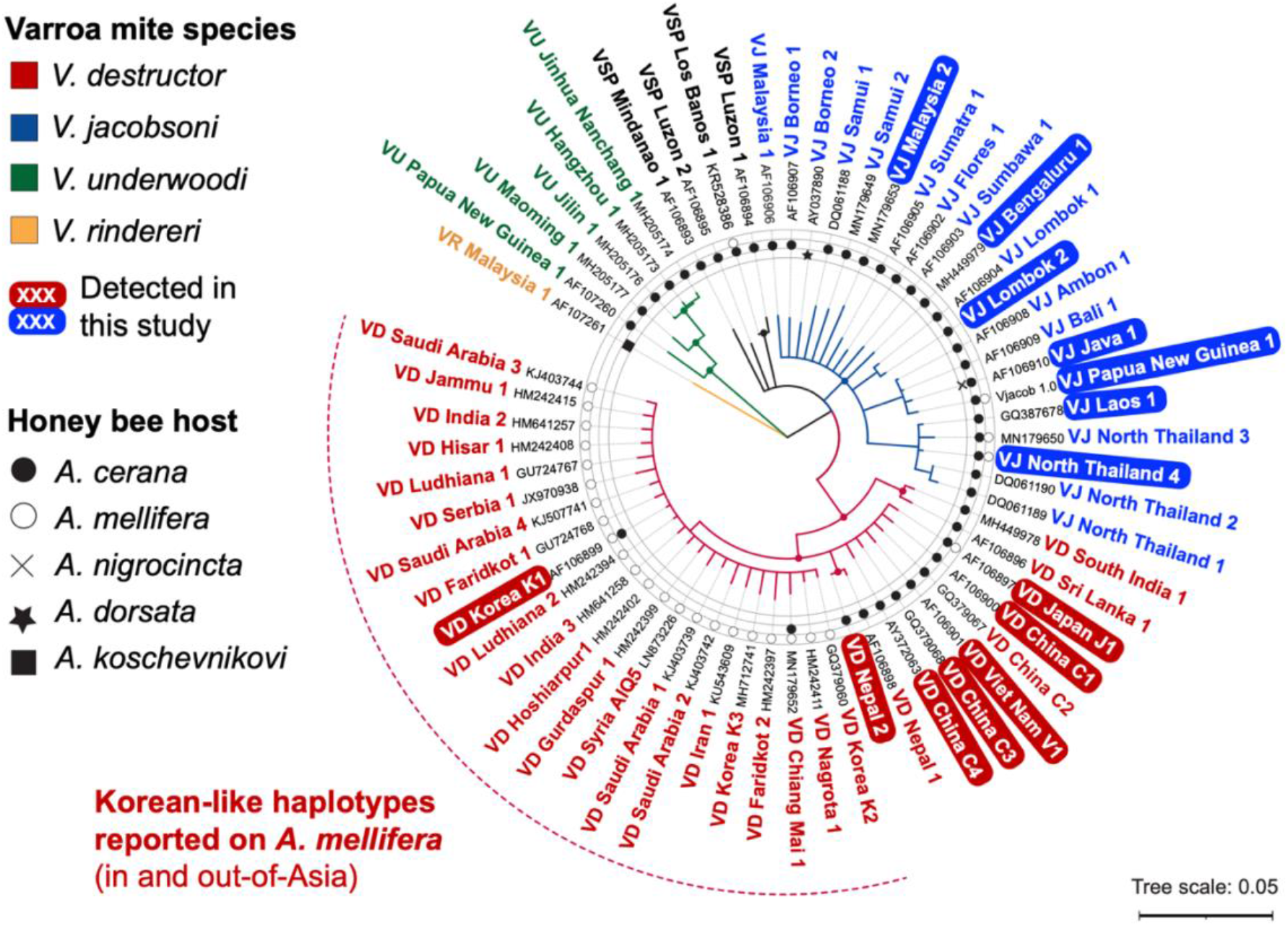
Sequenced *V. destructor* and *V. jacobsoni* mites originated from diverse maternal genetic backgrounds (lineages) shown by mtDNA *COX1* barcoding. A neighbor-joining tree was built using the Tamura-Nei genetic distance model with 1,000 bootstraps (> 80% are shown as filled circles in the base of each branch). *V. rindereri* was considered as an outgroup and the host range indicated [3]. *V. destructor* mites collected throughout Asia on their original host belong to seven distinct lineages (see details in Table S2). By contrast, all the host-switched mites across that range belong to the Korean K1 haplotype [AF106899], which is also found worldwide [5]. *V. jacobsoni* mites collected throughout Southeast Asia and Oceania, we identified seven lineages including two newly described: Lombok 2 and North Thailand 4. All mites parasitizing *A. mellifera* in Papua New Guinea were infested only by the *V. jacobsoni* Papua New Guinea 1 haplotype. The novel haplotype, North Thailand 4, was found on one *A. melli*fera colony in the northern limit of *V. jacobsoni* native range.

**Figure S2:**
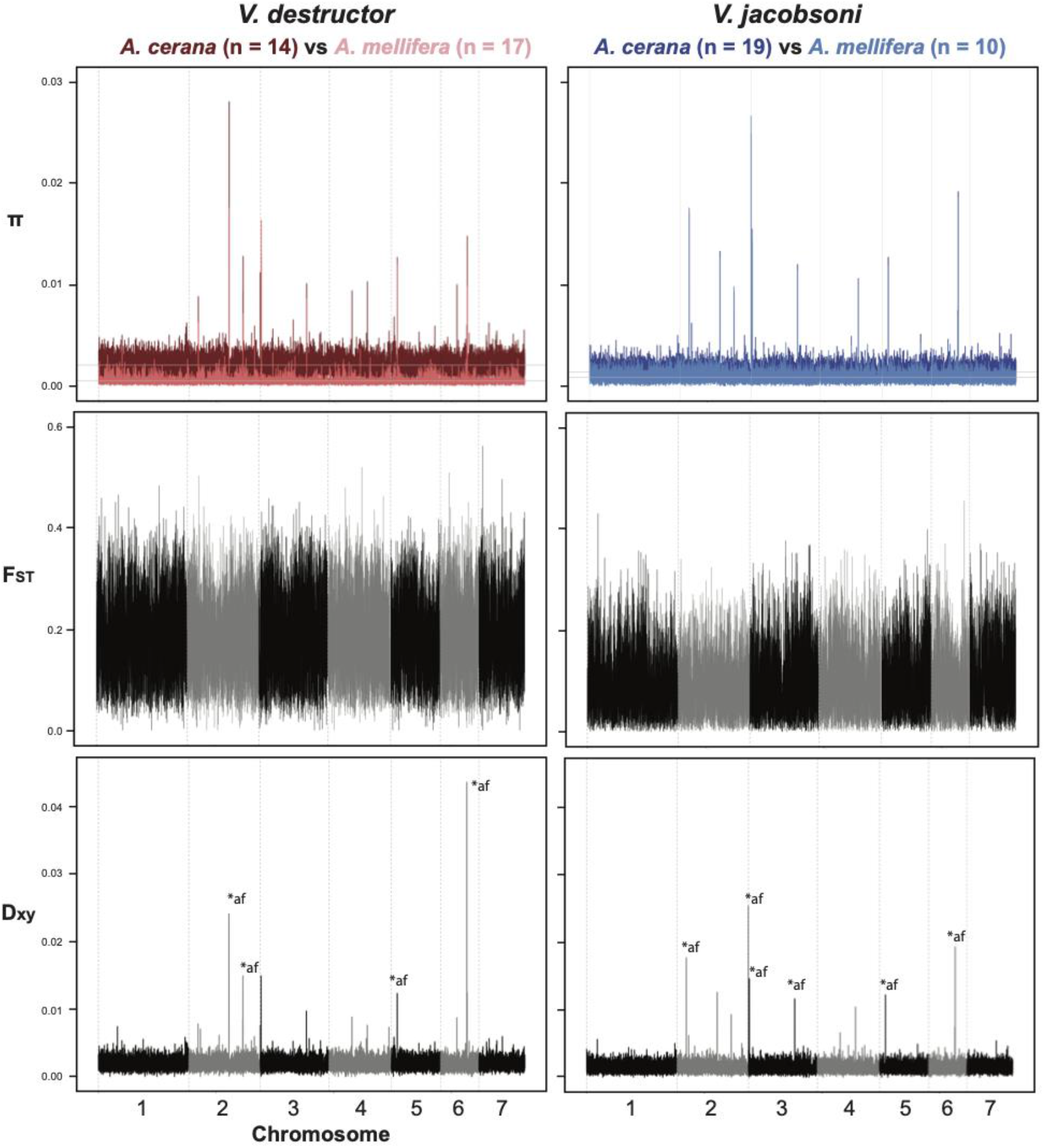
Parallel reduction of genetic diversity and genomic divergence between hosts in *V. destructor* and *V. jacobsoni*. First row: Nucleotide diversity in 20kb window along all seven chromosomes with mites parasitizing the same host pooled as one population for each *Varroa* species. Genetic divergence estimated by pairwise F_ST_ (second row) and absolute divergence using D_xy_ estimate (third row) in 20kb window, show a strong host differentiation all along the genome with few outliers. All sites with D_xy_ value > 0.01 were investigated manually by examining individual genotypes, which showed an excess in heterozygosity levels. These regions were also poorly conserved within species and had BLAST matches elsewhere in the genome, suggesting the presence of sequencing artefacts rather than biological signal (labelled *af on the plots).

**Figure S3.**
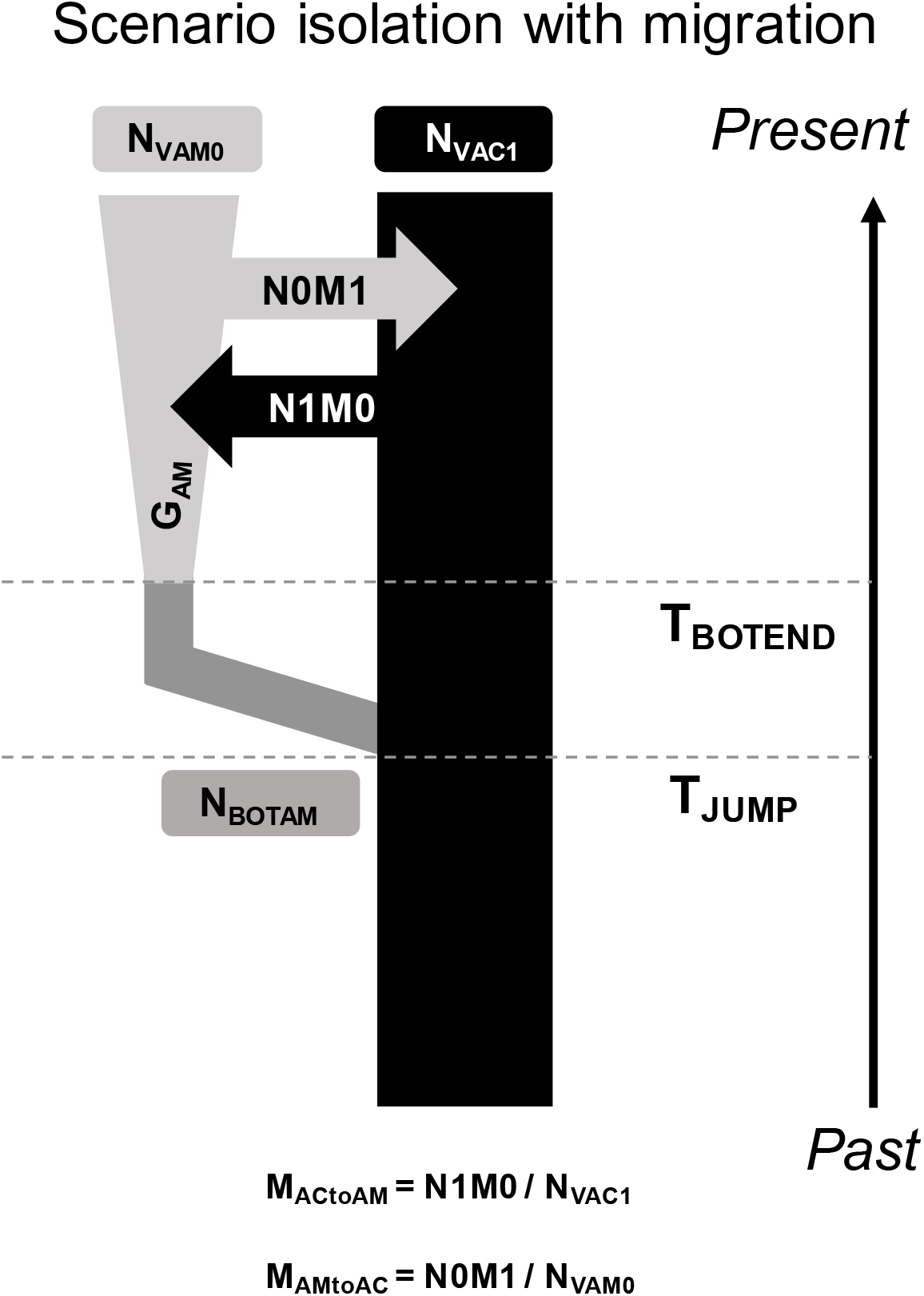
Graphical representation of *Varroa* host switch demographic scenario designed following population genomics analysis. The model was run independently for each *Varroa* species but contained the same parameters. N_VAM0_ is the present effective population size on the novel host (*A. mellifera*) whereas N_VAC1_ is the present population size on the original host (*A. cerana*). At a time in the past, T_JUMP_, a founding size of N_BOTAM_ switched durably host from *A. cerana* to *A. mellifera*. The founder effect associated with host switch ended at T_BOTEND_ and population on novel host expanded at a growth rate G_AM_. As both hosts are in artificial sympatry with evidence of gene flow, migration rates were estimated for spillover (M_ACtoAM_) and spillback (M_AMtoAC_). See Table 2 for parameters prior details.

**Figure S4.**
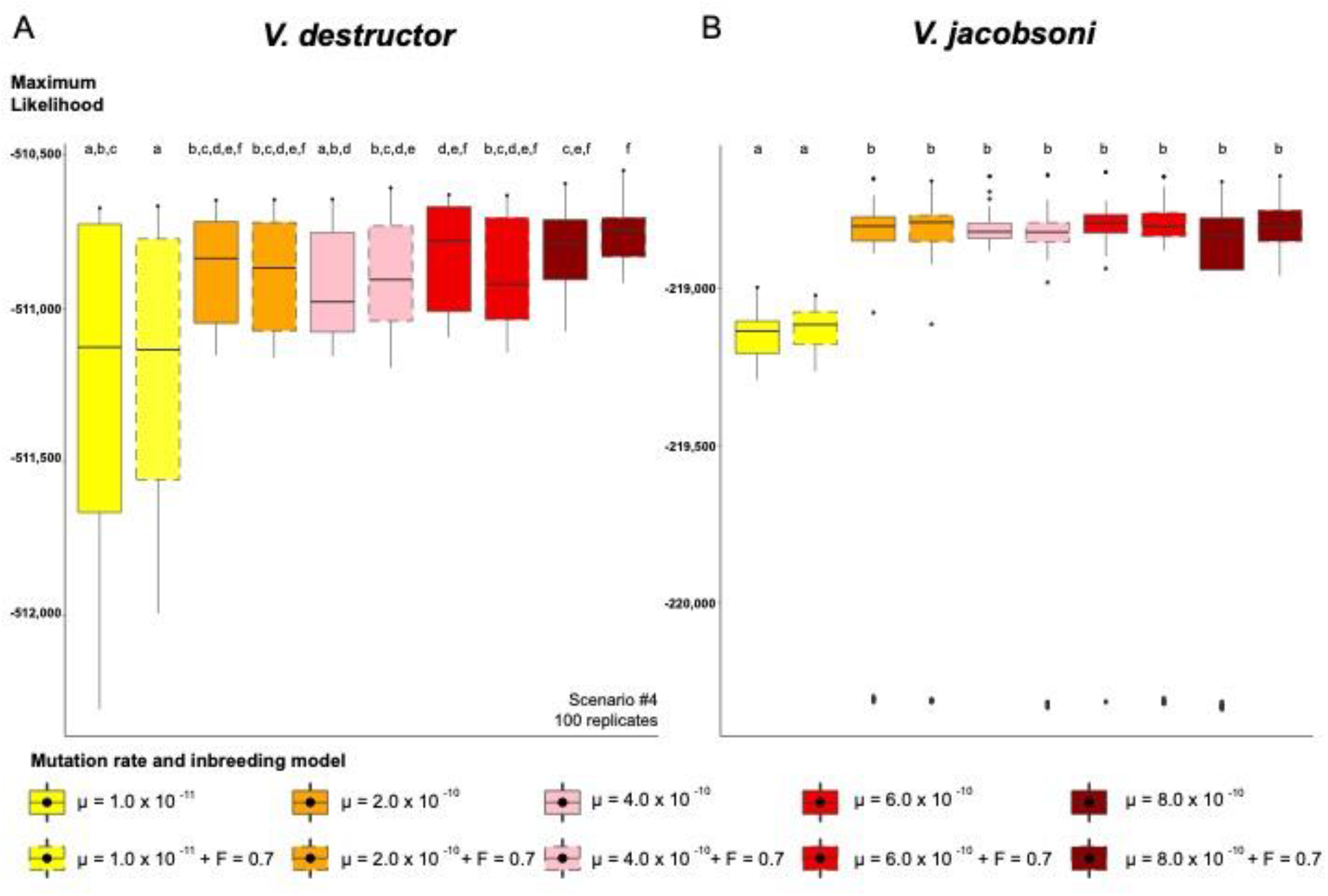
Distribution of the maximum likelihood obtained from the best pseudo-observed SFS generated by demographic inferences across different mutation rates and inbreeding levels in *Varroa* species. Replicates generated with lower mutation rates than the directly estimated 8.0 × 10^−10^ were not associated with a better fit to the observed SFS from our genomic data for *V. destructor* (A) and *V. jacobsoni* (B). No significant differences were detected in the likelihood variance among different mutation rates. However, pairwise Wilcoxon test significance (letters) after Bonferroni corrections showed that the lower mutation rate differentiated from other rates, especially in *V. jacobsoni*.

**Figure S5:**
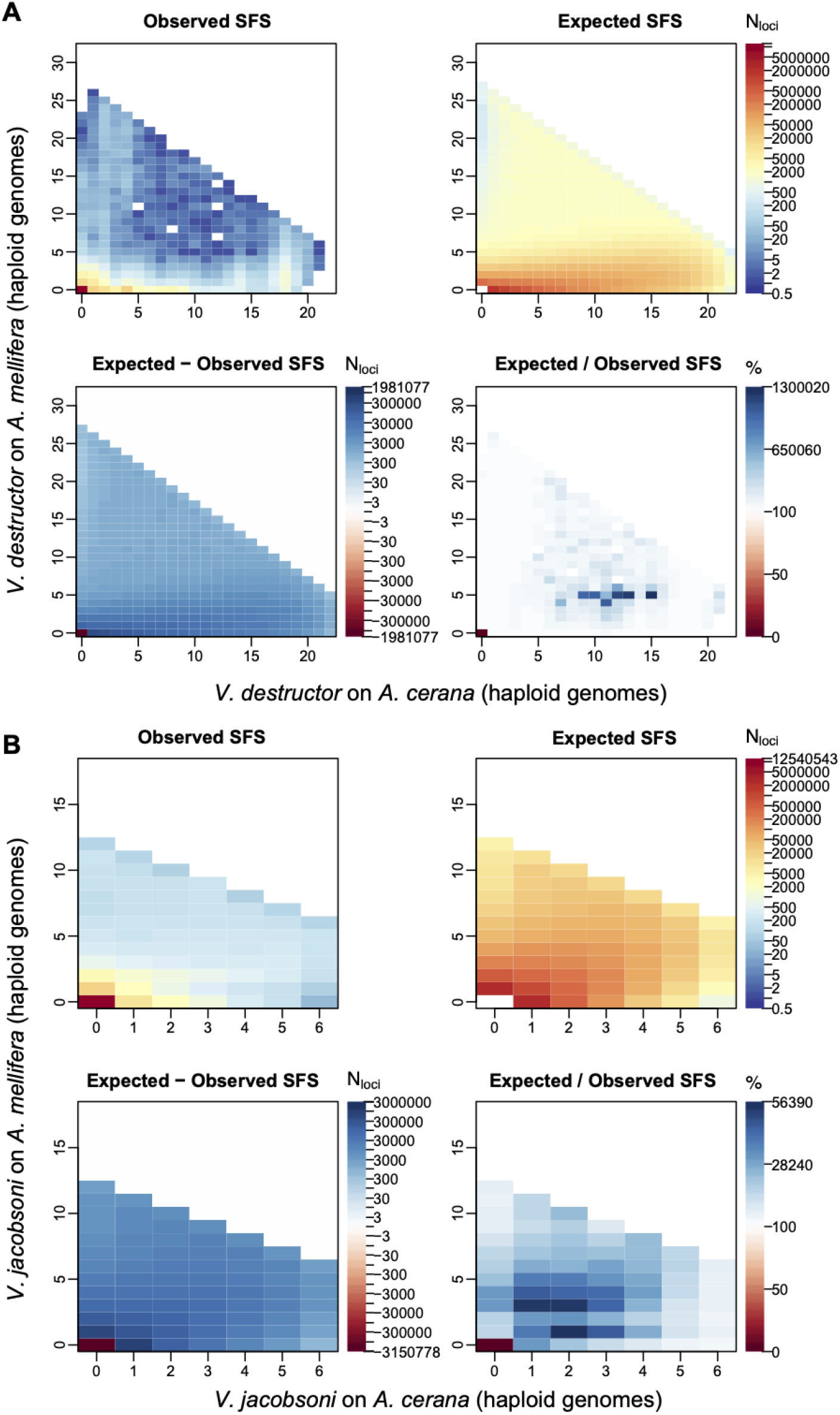
Observed vs expected joint 2D-SFS from the best replicate of the isolation with migration scenario for *V. destructor* and *V. jacobsoni*.

**Table S1: Detailed sampling information for each of the *Varroa* mite sequenced for population genomics as well as for estimation of the mutation rate.** Sampling details include *Varroa* species, host identity, country and locality, date and geographical coordinates. Sequencing details include the DRA accession on NCBI, the total number of reads, mapped reads, mapping ratio and mean read depth. Mitochondrial lineages and mitotype identity are also included for each specimen.

**Table S2: Repertoire of standard mtDNA markers used to determine mite species and lineages of sequenced specimens.** Names given to new mitotype were chosen to follow the revised nomenclature and compared to *V. destructor* and *V. jacobsoni* reference sequences (source and sequence accession indicated).

**Table S3: Observed (O_HOM_) and expected (E_HOM_) homozygosity and inbreeding coefficient (F) values computed for each sequenced mite on N_SITES_ for the population genomics analysis.**

**Table S4: Pairwise Weir and Cockerham F_ST_ values computed between host-adapted *Varroa* species and between host-adapted *Varroa* mtDNA lineages.** The color shades indicate relatively high genetic differentiation (in red), moderate (in yellow) to a low one (in green).

**Table S5: Number of reads from whole-body *Varroa* sample mapped against *A. mellifera* and *A. cerana* mitochondrial reference genomes.** Host metagenomics identity was assigned by the higher number of reads and compared to the field collection information. The proof of concept was confirmed with Mom and Son samples collected while feeding on their *A. mellifera* host.

